# POU6F2 Positive Retinal Ganglion Cells a Novel Group of ON-OFF Directionally Selective Subtypes in the Mouse Retina

**DOI:** 10.1101/2020.02.28.968503

**Authors:** Ying Li, Jiaxing Wang, Rebecca King, Eldon E. Geisert

## Abstract

**Purpose:** Previously we identified POU6F2 as a genetic link between central corneal thickness (CCT) and risk of open-angle glaucoma. The present study is designed to characterize the POU6F2-positive retinal ganglion cells (RGCs).

**Methods:** The Thy1-YFP-H mouse was used to identify the structure of POU6F2-positive RGCs in the retina. In the retina of the Thy1-YFP-H mouse approximately 3% of the RGCs were labeled with yellow fluorescent protein. These retinas were stained for POU6F2 to identify the morphology of the POU6F2 subtypes in 3D reconstructions of the labeled RGCs. Multiple retinal cell markers were also co-stained with POU6F2 to characterize the molecular signature of the POU6F2-positive RGCs. DBA/2J glaucoma models were used to test the role of POU6F2 in injury.

**Results:** In the retina POU6F2 labels 32.9% of the RGCs in the DBA/2J retina (16.1% heavily and 16.8% lightly labeled). In 3D constructions of Thy1-YFP-H positive RGCs, the heavily labeled POU6F2-positive cells had dendrites in the inner plexiform layer that were bistratified and appeared to be ON-OFF directionally selective cells. The lightly labeled POU6F2 RGCs displayed 3 different dendritic distributions, with dendrites in the ON sublaminae only, OFF sublaminae only, or bistratified. The POU6F2-positive cells partially co-stained with *Cdh6*. The POU6F2-positive cells do not co-stain with CART and SATB2 (markers for ON-OFF directionally selective RGC), SMI32 (a marker for alpha RGCs), or ChAT and GAD67(markers for amacrine cells). The POU6F2-positive cells were sensitive to injury. In DBA/2J glaucoma model, at 8 months of age there was a 22% loss of RGCs (labeled with RBPMS) while there was 73% loss of the heavily labeled POU6F2 RGCs.

**Conclusions:** POU6F2 is a marker for a novel group of RGC subtypes that are ON-OFF directionally selective RGCs that are sensitive to glaucomatous injury.

## INTRODUCTION

The present study focuses on POU6F2, a transcription factor marking a subset of retinal ganglion cells (RGCs) that are selectively sensitive to glaucomatous injury. Our interest in POU6F2 began with a series of studies using a mouse model system to define the genetic basis for phenotypic glaucoma risk factors. Glaucoma is a diverse set of diseases that if left untreated leads to permanent damage of axons in the optic nerve and visual field loss. Glaucoma affects millions of people worldwide ^1, 2^ and is the second leading cause of blindness in the United States ^3^. Adult-onset glaucoma is a complex collection of diseases with multiple risk factors and genes with differing magnitudes of effects on the final common pathway of glaucoma, the progressive loss of RGCs ^4–7^. The Ocular Hypertension Treatment Studies (OHTS) ^8^ and subsequent independent findings of others ^9, 10^, define the number of phenotypic risk factors for primary open angle glaucoma (POAG), including central corneal thickness (CCT). The thinner corneas are associated with an increased risk of developing POAG and this risk is independent of the confounding effects of CCT on intraocular pressure measurements ^8, 10^. The thinner CCT is also associated with an increased severity of visual field loss and a more rapid progression of the disease ^11–13^. We use a systems genomics approach (the BXD recombinant mouse strain and the powerful bioinformatics available on GeneNetwork.org) to define genomic elements modulating CCT ^14–19^. We examine the BXD strains and map genomic loci modulating CCT in the mouse ^15^. CCT was measured in 818 eyes in 61 BXD strains ^15^ and the data mapped to a single locus in the mouse. Within this quantitative trait loci (QTL) there is only one candidate gene in the mouse, *Pou6f2*. Our collaborator, Dr. Janey Wiggs examined these syntenic regions in the human NEIGHBORHOOD database ^20^ to determine if there are any potential risk factors for glaucoma within these loci. The top 50 hits are all one gene, *POU6F2* ^*15*^. At the time *POU6F2* was a potential risk factor for human glaucoma ^21, 22^; however a subsequent study from the UK Biobank ^23^ demonstrates that *POU6F2* is a glaucoma risk factor in humans even after correcting for multiple tests.

POU6F2 was first described as a novel POU-domain transcription factor in the retina^24^ and it identified a subpopulation of RGCs^24^. We have independently confirmed these findings and found that *Pou6f2* is part of a genetic network found in mouse RGCs ^25^. The first hint of the link between *Pou6f2* modulating CCT in the mouse and its potential role in glaucoma is revealed during the development of the eye. POU6F2 is expressed in the RGC progenitor cells and cornea. In the embryonic eye, strong POU6F2 staining was observed in neuroblasts destined to become RGCs. There is also staining of the developing cornea and corneal stem cells ^15^. The present study identifies multiple POU6F2-positive RGC subtypes, and a portion of these cells form a novel group of ON-OFF directionally selective RGCs.

## METHODS

### Mice

Thy1-YFP-H mice (B6.Cg-Tg(Thy1-YFP)HJrs/J) were purchased from Jackson Laboratory (Stock#:003782, Bar Harbor, ME, USA). In the Thy1-YFP-H mouse line approximately 3%∼10% of the RGCs are labeled by YFP in the retina^26^. This allows us to identify the morphology of RGCs that were positive for the marker POU6F2. We also stained retinas from *Cdh6*-CreER mice (B6.Cg-*Cdh6*^*tm1*.*1(cre/ERT2)Jrs*^/J, Jax Stock No:029428)^27^ to determine if there were independent markers for the POU6F2 cells identifying the specific POU6F2-positive ganglion cell subtype^28^. The DBA/2J mice were purchased from The Jackson Laboratories (Stock No:000671). To study the effects of glaucoma on retinal ganglion cell subtypes we used the DBA/2J model comparing aged DBA/2J mice (8 months old) to young adult DBA/2J mice (2 months old). We also studied the distribution of POU6F2-positive cells in C57BL/6J mice (n= 4 per group) and BALB/c mice (n= 4 per group).

All procedures involving animals were approved by the Animal Care and Use Committee of Emory University and were in accordance with the ARVO Statement for the Use of Animals in Ophthalmic and Vision Research. The mice were housed in a pathogen-free facility at Emory University, maintained on a 12 h:12 h light–dark cycle, and provided with food and water ad libitum.

### Immunohistochemistry

For immunohistochemistry, the mice were deeply anesthetized with ketamine 100mg/kg and xylazine 15 mg/kg and perfused through the heart with saline followed by 4% paraformaldehyde in phosphate buffer (pH 7.3). The eyes were removed, and the retinas were dissected out. Flat-mounts of the retina were prepared for staining using a protocol similar to that previously described ^15^. Briefly, the retinas were stained for POU6F2 with a rabbit antibody against POU6F2 (MyBiosource, Cat. # MBS9402684, San Diego) at 1:500 and to label POU6F2-positive RGCs. Other antibodies used in this study are listed in Table 1. After the primary antibodies staining for over two nights, the retinas were rinsed and placed in secondary antibodies which included Alexa Fluor 488 AffiniPure Donkey Anti-Rabbit, Cat. #715-545-152 or Alexa Fluor 594 AffiniPure Donkey Anti-Guinea pig, Cat. #706-585-148, Jackson Immunoresearch, West Grove). For molecular characterization of the POU6F2-positive RGCs, the retinal flat-mounts were also stained for TO-PRO-3 to label nuclei in the retina. For morphological characterization of the POU6F2-positive RGCs, the retinas were counterstained for goat anti-ChAT (Table 1) to label ChAT-positive amacrine cell dendrites as a marker for sublaminae S2 and S4 of the inner plexiform layer ^29^.

**Table 1.**
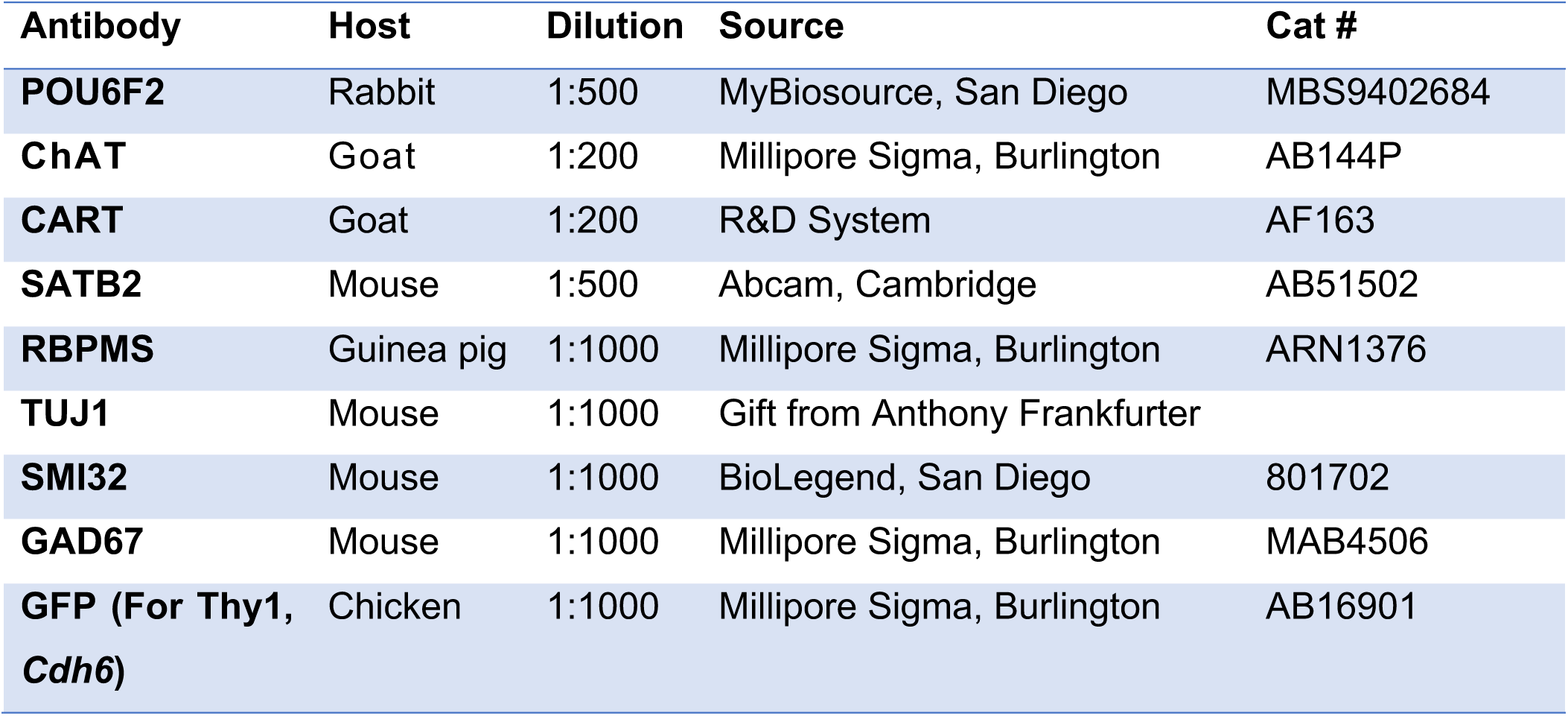
Antibodies used in this study.

### Image processing and statistics

All retinal flat-mounts were imaged using a Nikon Eclipse Ti (Nikon, Inc., Melville, NY, United States) confocal microscope. To identify the specific morphology of the POU6F2 RGCs, the double labeled RGCs (POU6F2-positive/GFP-positive) in Thy1-YHP-H retinal flat-mounts were identified and scanned under 40X magnification including all dendrites. Z-stacked images were taken at 0.1 μm increments, with a total of 300–600 optical slices for each retina. Later, 3D reconstructions of the POU6F2-positive cells were made from z-stacks of confocal images using the Imaris (v9.2, Bitplane, Oxford Instruments, Switzerland). The 3D reconstructions can be rotated 90° to identify the sublaminae distribution of the ganglion cell dendrites (Figure 1).

**Figure 1.**
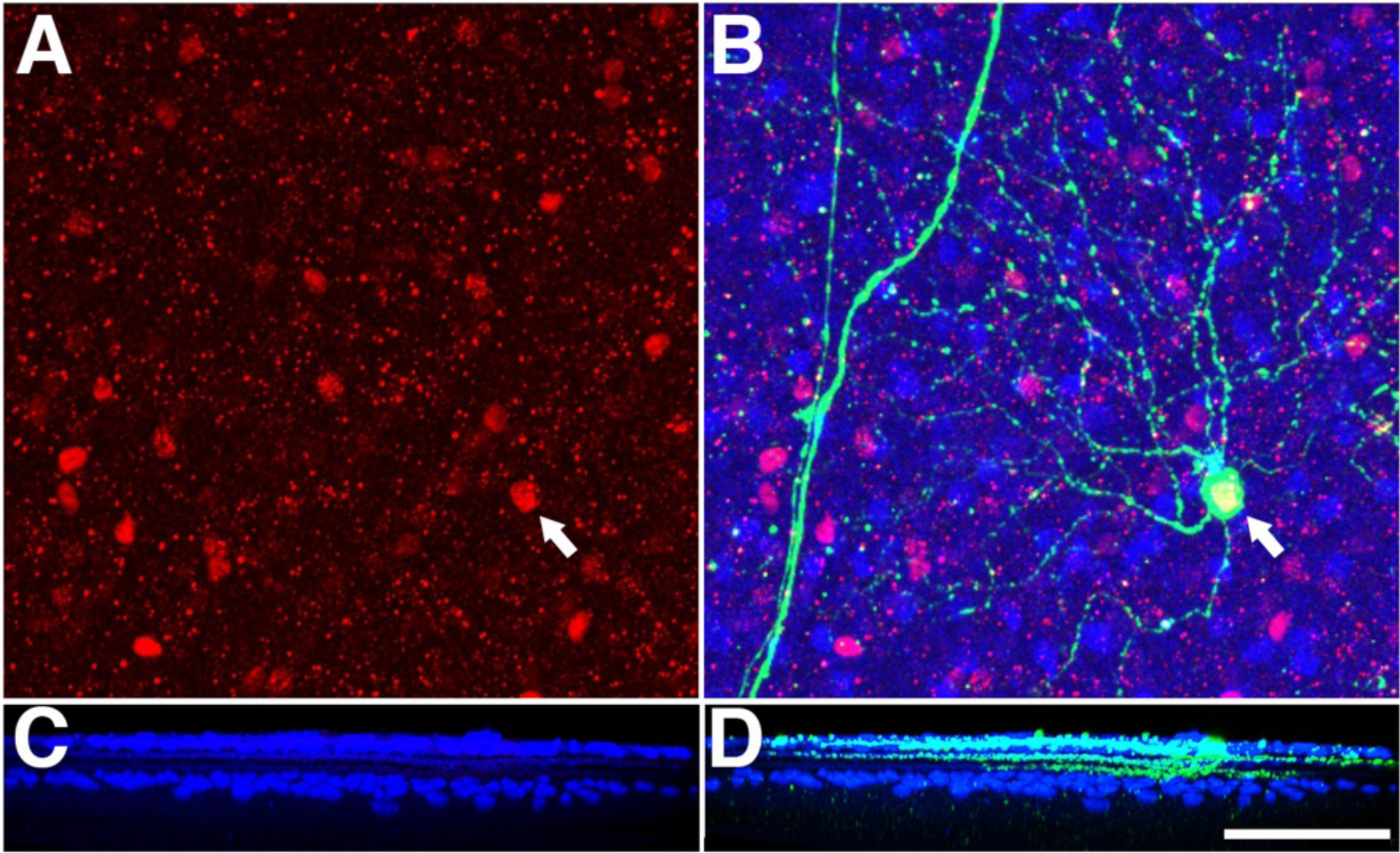
Dendritic morphology of heavily labeled POU6F2 cells (A) in retina of Thy1-YFP-H mice. In the merged channel (B) the POU6F2-positive nucleus (Red) can be seen in the YFP labeled RGC (Green) marked by the arrow. Amacrine cells labeled with ChAT (Blue) mark the location of sublaminae S2 and S4(C). A 90 degree rotation of the 3D reconstruction is shown in D. Notice the distribution of labeled dendrites in the ON and OFF sublaminae showing the POU6F2 RGC is bistratified. Scale bar equals 100µm.

To identify the molecular characteristics of POU6F2-positive RGCs, the density of RGCs was calculated according to the number of labeled cells per field within the ganglion cell layer using CellProfiler (v3.1.5) ^30, 31^. The pipeline follows the method from Dordea, et al ^32^ with modifications to optimize the counting for POU6F2 and RPBMS positive cells. For each RGC marker, 4 independent biological retina samples were examined for the quantification of RGCs per field. For each retina, four fields with equal distance to the optic nerve were sampled from each quadrant of the middle regions of each retina under 20x magnification.

Data are presented as mean ± standard error of the mean (SEM). Differences in RGC counts were analyzed with the Mann–Whitney U-test using SPSS statistics package 24.0 (SPSS, IBM, Chicago, IL, United States). A *p*-value of less than 0.05 was considered statistically significant.

## RESULTS

### Morphological Characterization of POU6F2 RGCs

In flat-mounts of the C57BL/6J mouse retina POU6F2 labels nuclei of cells that were also positive for RBPMS (a pan-RGC marker). The intensity of the POU6F2 labeling varied considerably. Some RGCs have highly fluorescent nuclei while others have lightly labeled nuclei. In the retinal ganglion cell layer of the C57BL/6J mouse, approximately 17.4% of the RBPMS-positive RGCs were heavily labeled with POU6F2. The remaining POU6F2-positive cells were moderately to lightly labeled and represented 18.1% of the RGCs. There was a modest number of POU6F2-positive cells in the inner surface of the inner nuclear layer, the amacrine cell layer. The POU6F2-positive cells at the surface of the inner nuclear layer were also positive for the RGC marker RBPMS, indicating that these cells may be displaced RGCs. To determine if the POU6F2 cells were all RGCs, we crushed the optic nerve and examined retinas 28 days after crush. All of the nuclear staining in the ganglion cell layer and in the amacrine cell layer was lost. These data support the conclusion that all of the POU6F2 positive cells are RGCs^15^.

The dendritic morphology of the POU6F2-positive cells was identified by staining the retinas of Thy-1-YFP-H mice with antiserum directed against POU6F2. Approximately 3% of the RGCs in the Thy-1-YFP-H mice are labeled with YFP, revealing the morphology of the cells including the dendritic arbors. In flat-mounts, YFP labeled RGCs co-stained with POU6F2 were identified and imaged to create a high-density z-stack. The retinas were counterstained for ChAT to label ChAT-positive amacrine dendrites as a marker for sublaminae S2 and S4 of the inner plexiform layer. 3D reconstructions of the POU6F2-positive cells were constructed using the Imaris software. The 3D reconstructions were then rotated 90° to identify the distribution of the ganglion cell dendrites in the inner plexiform layer (Figure 1). This allowed us to identify individual POU6F2 cells based on the dendritic morphology as well as the distribution of dendrites within the inner plexiform layer. A total of 12 heavily labeled POU6F2 RGCs and 4 lightly labeled POU6F2 RGCs were examined. Of the 12 heavily labeled RGCs, all had dendrites that were bistratified in the inner plexiform layer, occupying sublaminae S2 and S4. Based on their dendritic morphology all of these cells were ON-OFF RGCs.

Furthermore, their dendritic expanse over the surface of the retina was consistent with the heavily labeled POU6F2 cells being directionally selective RGCs. The lightly labeled cells displayed a different laminar distribution. The cells had dendrites ramifying in the ON sublaminae of the inner plexiform layer (n=1) or in the OFF sublaminae (n=2) or in the ON-OFF sublaminae (n=1).

### Molecular Characterization of POU6F2 RGCs

To aid in characterizing the POU6F2 RGCs, we stained the flat-mounts of the retina for POU6F2 plus a second RGC subtype specific marker. Retinas stained with secondary antibody only were used as controls. The markers we used are listed in Table 2. We first co-stained POU6F2 with pan-RGC markers including RBPMS, TUJ1 and Thy1 and found that all POU6F2-positive cells are co-stained with the pan-RGC markers, confirming that in the adult mouse retina, all the POU6F2-positive cells were RGCs. Since the heavily labeled POU6F2 cells were all ON-OFF RGCs based on the morphological findings, we examined other markers ON-OFF directionally selective RGCs. The first marker we co-stained with was Cocaine- and Amphetamine-regulated transcript (CART) a marker for most known ON-OFF directionally selective RGCs in the mouse ^33–36^. None of the POU6F2 cells were positive for CART. This was the first evidence that the POU6F2 cells may represent a novel subclass of RGC. We also stained for retinas for SATB2 a known marker of ON-OFF directionally selective RGCs^25, 37, 38^ and again none of the POU6F2-positive cells were positive for SATB2 (Table 2). We used a reporter mouse (*Cdh6*-CreER) ^33, 36^ to label *Cdh6-*positive cells with GFP. The retinas were stained for POU6F2. There were cells in the retina that were POU6F2-positive RGCs and also labeled with *Cdh6* driven GFP (Figure 2). Results showed that in double-labeled retinas, about 50% of the *Cdh6*-positive cells are also positive for POU6F2 (Figure 2). For the heavily labeled POU6F2-positive cells (16% of the total RGCs) approximately 25% are *Cdh6* positive (4% of the total retinal RGCs), The remaining 75% are *Cdh6* negative (12% of the total RGCs). Beyond the ON-OFF RGC markers, we also co-stained retinas for POU6F2 with other cell markers such as SMI32 (Alpha-RGC marker), GAD67 and ChAT (Amacrine cell markers). None of these markers are shown to co-localize with POU6F2 (Table 2).

**Table 2.** Co-staining of RGC markers with POU6F2.

| Marker | Marker | Strain | Co-staining * |
| --- | --- | --- | --- |
| Thy1 | Pan-RGCs | Thy1-CFP reporter mice | 100% |
| RBPMS | Pan-RGCs | C57BL/6J | 100% |
| TUJ1 | Pan-RGCs | C57BL/6J | 100% |
| *Cdh6* | On-Off DSGCs | *Cdh6*-CreER | 50% |
| CART | On-Off DSGCs | C57BL/6J | No |
| SATB2 | On-Off DSGCs | C57BL/6J | No |
| SMI32 | alpha RGCs | C57BL/6J | No |
| ChAT | Amacrine | C57BL/6J | No |
| GAD67 | Amacrine | C57BL/6J | No |

\*Co-staining represents the percentage of POU6F2-positive cells in the total population of the marker-positive cells.

**Figure 2.**
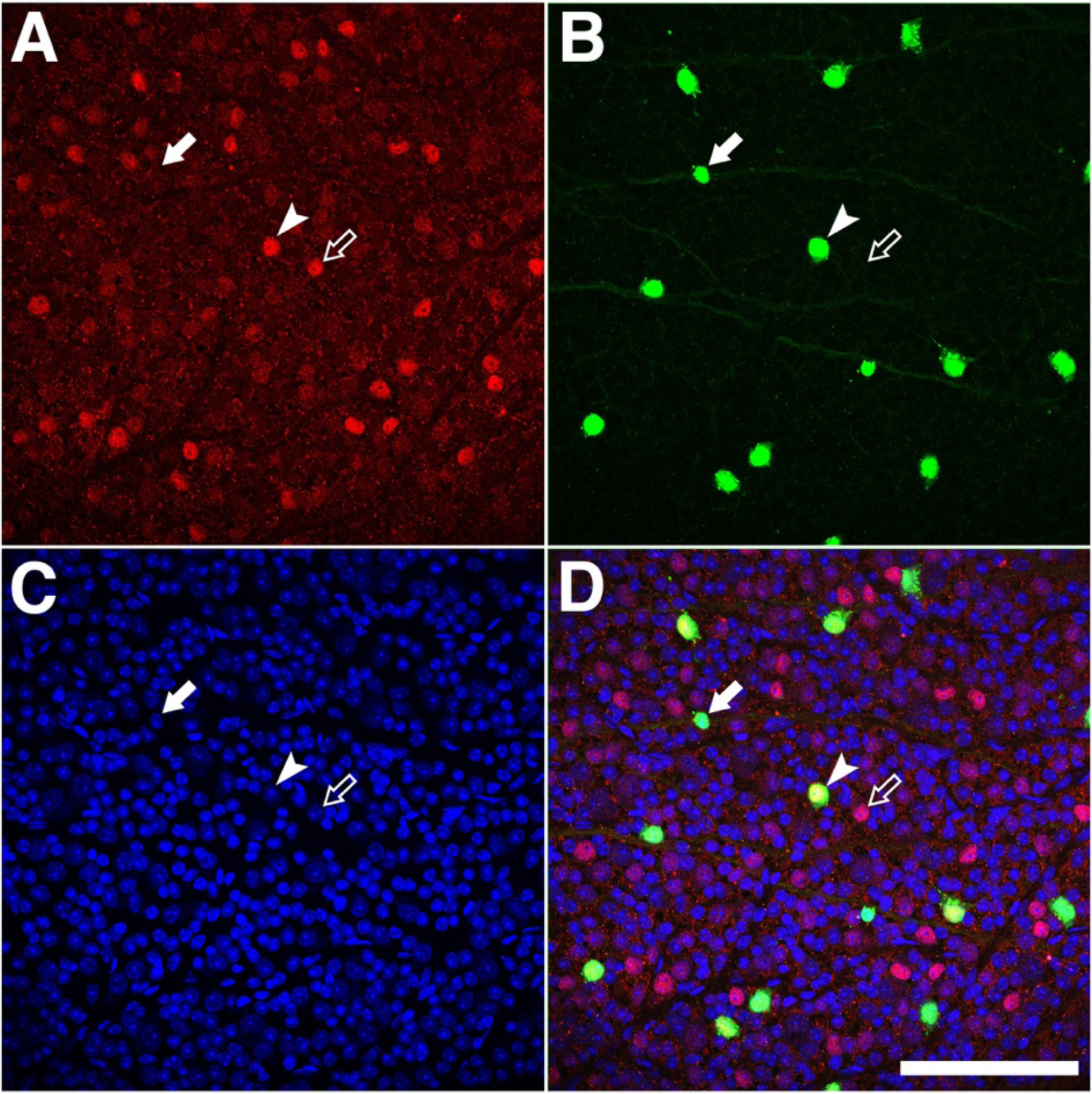
Retinal flat mounts of the *Cdh6*-CreER mice were stained for POU6F2 (A). The labeling of *Cdh6* is shown in B and TO-PRO-3 nuclear labeling is shown in C. The merged image is shown in (D). RGCs are present that are labeled by POU6F2 only (Empty Arrows), and *Cdh6* only (Filled Arrows). Approximately half of the *Cdh6*-positive cells were also positive for POU6F2 (Arrowhead, Double labeled cell). The scale bar in D equals 100µm.

### Mouse Strain Differences in POU6F2 RGCs

We examined the distribution of POU6F2 positive RGCs in three strains of mice commonly used in vision research: C57BL/6J, DBA/2J and BALB/c. In flat-mounts of the retinas the cells were stained for POU6F2 and RBPMS (Figure 3). This analysis revealed a significant difference in the number of POU6F2 positive RGCs in the different strains. The adult C57BL/6J retina had a total of 35.5% of the RGCs being POU6F2 positive with 17.4% heavily labeled and 18.1% lightly labeled. A similar density of POU6F2 positive cells were observed in the DBA/2J mouse retina, revealing a total of 32.9% positive for POU6F2, with 16.1% being heavily labeled and 16.8% being lightly labeled. The examination of the BALB/c retina showed a significantly higher number of RGCs labeled with POU6F2 (Figure 3). In the BALB/c retina there was a total of 65% of the RGCs labeled with POU6F2, with 27% being heavily labeled and 37% of the cells were lightly labeled. In all three strains all of the POU6F2 cells were also labeled with RBPMS in the adult, indicating that POU6F2 only labels RGCs in the adult retina.

**Figure 3:**
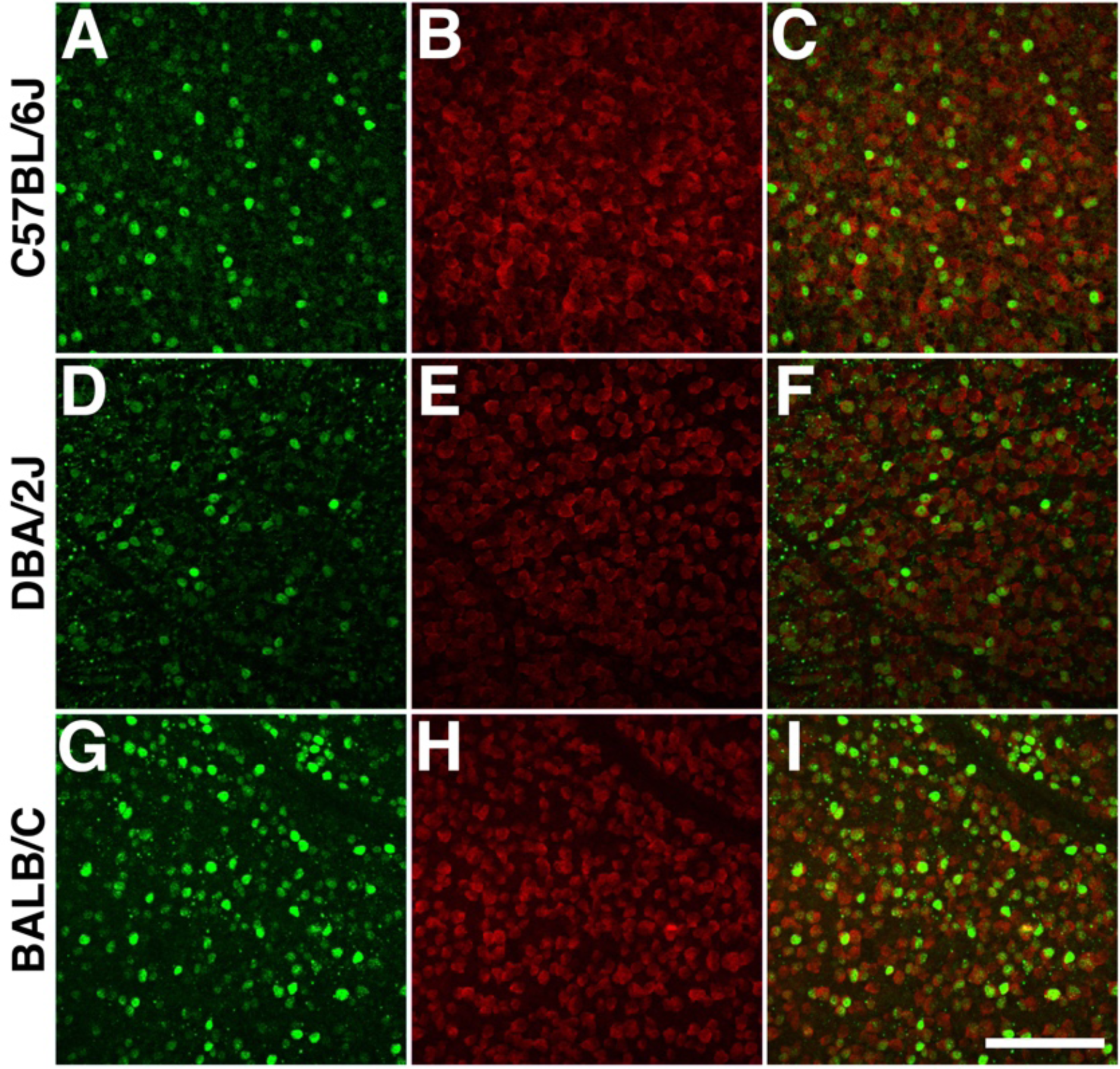
Strain differences in POU6F2 positive RGCs. RGCs were stained for POU6F2 (Green) and RBPMS (Red) in retinal flat mounts in 3 different strains of mice: C57BL/6J (A-C), DBA/2J (D-F) and BALB/c (G-I). Notice that the number of POU6F2 positive cells are approximately equal in the C57BL/6J (A-C), DBA/2J (D-F) retinas; while there are significantly more in the BALB/c retina. Scale bar in I equals 100µm.

### Effects of Glaucoma (DBA/2J Mouse Model)

In our previous study, we reported that the heavily labeled POU6F2 RGC subtype was very sensitive to early phases of glaucoma in the DBA/2J model ^15^. This analysis was expanded to examine the distribution of heavily labeled POU6F2-positive RGCs and lightly labeled POU6F2 RGCs. There was a 22% loss of RPBMS labeled RGCs in the aged mice in the aged D2 mice and a 73% loss of POU6F2 heavily labeled RGCs. When we examined the lightly labeled POU62 RGCs there was only 10% loss of these cells (Figure 4). If we examine the loss of RGCS labeled with RBPMS that are not heavily labeled with POU6F2, then there is an 11% loss of RBPMS-positive /POU6F2-negative RGCs. This loss of RPBMS-positive – heavily labeled POU6F2 RGCs (11%) is approximately equal to the loss of lightly labeled POU6F2-labeled RGCs (10%) (Figure 4). These data demonstrate that the heavily labelled POU6F2 RGCs were lost in the early phases of glaucoma in the DBA/2J mouse.

**Figure 4.**
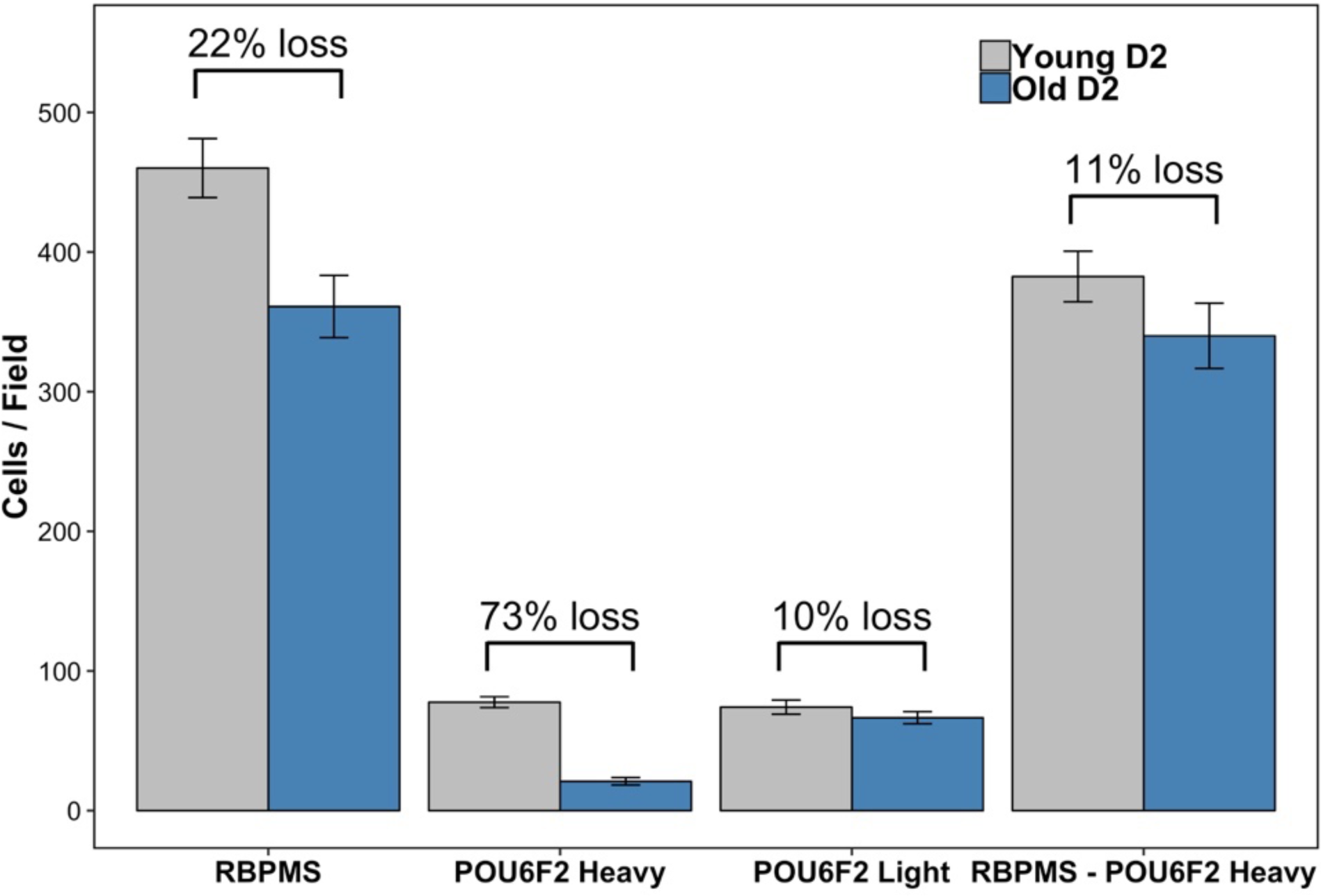
The selective sensitivity of POU6F2 RGC subtypes was demonstrated using the DBA/2J model of glaucoma. There was a 22% loss of RBPMS-labeled RGCs in aged DBA/2J mice (8 months of age, Old D2) as compared to young DBA/2J mice (2 months of age, Young D2). There was a dramatic loss (73%) of heavily labeled POU6F2-positive cells and was a mild loss (10%) of the lightly labeled POU6F2 RGCs in the Old D2 mice when compared with Young D2. When we exclude the heavily labeled POU6F2-positive cells, there is an 11% loss of the rest cells, approximately the same percentage of cell loss when compared with the lightly labeled POU6F2-positive RGCs. These data demonstrate the sensitivity of the POU6F2 RGC subtypes to glaucoma.

## DISCUSSION

We examined the distribution of POU6F2 in flat mounts of the DBA/2J and C57BL/6J retina. In the adult retina, all of the POU6F2-positive cells were also labeled with RBPMS, indicating that these cells are RGCs ^15^. The intensity of POU6F2 labeling varied considerably. We divided the cells into heavily labeled POU6F2 RGCs and lightly labeled POU6F2 RGCs. The lightly labeled cells had less than 50% of the fluorescent intensity as do the heavily labeled RGCs. The heavily labeled POU6F2 RGCs made up approximately 16.8 % of RPBMS RGCs in the DBA/2J retina and lightly labeled RGCs were approximately 16.1 % of the RPBMS RGCs. A few cells in the amacrine cell layer were also positive for POU6F2. All of these cells in the amacrine layer were also labeled with RBPMS, suggesting that these cells were in fact displaced ganglion cells. In retinas 28 days following optic nerve crush no RGCs in the retina were positive for POU6F2. Taken together, these data indicate that POU6F2 only labeled RGCs in the adult DBA/2J mouse retinas and similar finding were observed in the C57BL/6J retina. We also found that there was a surprising difference in the number of POU6F2-positive cells between the BALB/c retina and the retinas of the DBA/2J mouse and C57BL/6J mouse, explaining in part the differences observed in our previous study ^15^ and the initial study from the Jeremy Nathans laboratory ^24^.

The classification of RGC subtypes is undergoing a revolution due to the unbiased quantitative approach of single cell RNA seq and transcriptional profiling of individual RGCs. The resultant clustering of the cells based on similar gene expression profiles clearly reveals RGC subtypes. The advent of single cell RNA seq protocols has opened this approach to study of the retina ^39, 40^ and retinal ganglion cells ^41, 42^. Two studies have characterized retinal ganglion cells from the C57BL/6J mouse: isolated at postnatal day 5 ^41^ or isolated from the adult mouse retina ^42^. When comparing the expression of *Pou6f2* across these studies the expression profiles within the population of RGCs differs significantly. To determine if this difference reflected the biology of the retina or technical differences, we stained retinas for POU6F2 at these different ages. We found that at P5, 51% of the RBPMS-positive RGCs were positive for POU6F2 and there was a small population that were POU6F2-positive and RBPMS-negative, presumptive amacrine cells (unpublished observation). In the present study we examined POU6F2 in the adult retina of three different mouse strains, finding that both C57BL/6J and DBA/2J mice had approximately the same number of POU6F2-positive RGCs, 35.5% for C57BL/6J and 32.9% for DBA/2J. The BALB/c mouse had significantly more POU6F2 positive cells, with 64% of the RGCs being positive for POU6F2. To compare our results to the single cell RNA seq data, we examined the adult data from Tran et al ^42^ in relation to our results from the adult C57BL/6J mouse retina. In the Tran et al. ^42^ dataset there are six RGC subtypes expressing relatively high levels of *Pou6f2* and all six of these were classified as Novel (7-Novel, 8-Novel, 10-Novel, 18-Novel, 37-Novel and 44-Novel). Similar to our classification based in reporter mice and immunostaining, three of these subtypes express *Cdh6* (7-Novel, 8-Novel and 10-Novel), while three subtypes do not (18-Novel, 37-Novel and 44-Novel). The total percentage of highly expressing *Pou6f2* cells is 15.9%, similar to our immunohistochemical findings. Finally, none of these *Pou6f2* expressing subtypes express CART. These findings from the Tran et al. ^42^ study confirm our observations in the C57BL/6J retina.

### Morphology of Heavily Labeled POU6F2 RGCs

To define the morphology of the POU6F2 labeled cells we examined the retinas of Thy1-YFP-H mice immunostained for POU6F2. All of the heavily labeled POU6F2 RGCs had dendrites in the sublaminae S2 and S4 indicating they were ON-OFF cells and their dendritic morphology was consistent with them being directionally selective. To identify the 12 RGCs presented in this paper we had to examine 10 flat mount retinas. In each retina there were approximately 200 RGCs labeled with YFP and of these on average only one cell was found co-stained for POU6F2 in each retina. If these numbers were taken in isolation one would believe that only 0.5% of the RGCs were heavily labeled POU6F2 RGCs. However, we know based on immunohistochemical staining of retina and data from single cell RNA seq studies ^42^ that the number of heavily labeled cells is approximately 16% of the total RGCs. Thus, it appears that the labeling in the Thy1-YFP-H mouse is not random but has an inherent bias, and in our case against POU6F2 RGCs. To further illustrate the bias in the Thy1-YFP-H mouse retina we looked at a study by Bray et al. study ^43^. These authors found that the majority of the labeled RGCs of the Thy1-YFP-H mice were αRGCs (OSPN-positive, ∼70%), while in single cell RNA seq studies ^42^ only approximately 8 % of the RGCs had an alpha RGC profile. Another example of potential bias comes from a study of directionally selective RGCs ^33^ where the authors found that all of the ON-OFF cells were CART positive and represented 15% of the total RGCs. We have not observed double labeling of RGCs with CART and POU6F2 suggesting they are potentially two separate populations. We know that on the basis of RNA profiling ^42^ these cells cell types represent multiple subclasses of RGCs. It is possible that specific subclasses of POU6F2 cells were not identified in the Thy1-YFP-H retina. Nonetheless, of the 12 heavily labeled POU6F2 cells reconstructed in this study all were ON-OFF directionally selective RGCs.

### POU6F2 RGCs susceptible to injury

The present study reveals that the heavily labeled POU6F2 cells are some of the first cells to go missing in the DBA/2J model of glaucoma. Based on our analysis of dendritic morphology these cells are ON-OFF directionally selective RGCs. Previous studies have indicated that the ON-OFF directionally selective RGCs were very susceptible to injury following optic nerve crush ^44 45^. In these studies, it was thought that the ON-OFF directionally selective RGCs represented approximately 15% of the total RGCs in the retina and all of these cells were CART positive ^33, 36, 44^. Based on our findings and those of Tran et al. ^42^ there is a need to reassess the classification of RGCs in the mouse as well as the susceptibility of these cells to glaucomatous insult and optic nerve crush. In this paper we demonstrate that 16% of the RGCs are POU6F2-positive, ON-OFF directionally selective RGCs and these cells are not CART positive. Previous studies ^33, 44^ found that 15% of the RGCs are CART positive, ON-OFF directionally selective RGCs. Interestingly, the most recent single cell RNA-seq study of the RGCs ^42^ defined transcriptome changes of RGCs in response to injury, optic nerve crush. Their data showed that *Pou6f2* is one of the genes that is sensitive to injury and is down regulated within 12 hours after optic nerve crush ^42^. These findings supported our observation that the POU6F2-positive cells are sensitive to glaucomatous injury in the DBA/2J mouse model of glaucoma ^15^.

We have identified a novel class of ON-OFF directionally selective RGCs heavily labeled by POU6F2. These cells are distinct from the CART positive ON-OFF directionally selective RGCs previously described ^33^. Thus, there are two distinct populations of ON-OFF directionally selective RGCs in the mouse retina, one POU6F2 positive and the other CART positive, making the total number of ON-OFF directionally selective RGCs close to 30% of the total mouse RGCs. At the present time it appears that the POU6F2 ON-OFF directionally selective cells are the most sensitive to glaucomatous injury. This sensitivity may be restricted to only a few subtypes of the heavily labeled POU6F2 cells.

## Acknowledgements

This study was supported by an Unrestricted Grand from Research to Prevent Blindness, Owens Family Glaucoma Research Fund (EEG) and National Eye Institute Grant P30EY006360 (Emory Vision Core). The granting agencies of this research have no role in study design, data collection and analysis, decisions to publish or preparation of the manuscript.

## REFERENCES

1. Quigley HA. Number of people with glaucoma worldwide. Br J Ophthalmol 1996;80(5):389–93.

2. Thylefors B, Negrel AD. The global impact of glaucoma. Bull World Health Organ 1994;72(3):323–6.

3. Leske MC. The epidemiology of open-angle glaucoma: a review. Am J Epidemiol 1983;118(2):166–91.

4. Liu Y, Allingham RR. Molecular genetics in glaucoma. Exp Eye Res 2011;93(4):331–9.

5. Aboobakar IF, Johnson WM, Stamer WD, et al. Major review: Exfoliation syndrome; advances in disease genetics, molecular biology, and epidemiology. Exp Eye Res 2016;154:88–103.

6. Springelkamp H, Iglesias AI, Mishra A, et al. New insights into the genetics of primary open-angle glaucoma based on meta-analyses of intraocular pressure and optic disc characteristics. Hum Mol Genet 2017.

7. Nickells RW. The cell and molecular biology of glaucoma: mechanisms of retinal ganglion cell death. Invest Ophthalmol Vis Sci 2012;53(5):2476–81.

8. Gordon MO, Beiser JA, Brandt JD, et al. The Ocular Hypertension Treatment Study: baseline factors that predict the onset of primary open-angle glaucoma. Arch Ophthalmol 2002;120(6):714-20; discussion 829-30.

9. Medeiros FA, Sample PA, Weinreb RN. Corneal thickness measurements and visual function abnormalities in ocular hypertensive patients. Am J Ophthalmol 2003;135(2):131–7.

10. European Glaucoma Prevention Study G, Miglior S, Pfeiffer N, et al. Predictive factors for open-angle glaucoma among patients with ocular hypertension in the European Glaucoma Prevention Study. Ophthalmology 2007;114(1):3–9.

11. Leske MC, Heijl A, Hyman L, et al. Predictors of long-term progression in the early manifest glaucoma trial. Ophthalmology 2007;114(11):1965–72.

12. Medeiros FA, Sample PA, Zangwill LM, et al. Corneal thickness as a risk factor for visual field loss in patients with preperimetric glaucomatous optic neuropathy. Am J Ophthalmol 2003;136(5):805–13.

13. Herndon LW, Weizer JS, Stinnett SS. Central corneal thickness as a risk factor for advanced glaucoma damage. Arch Ophthalmol 2004;122(1):17–21.

14. Geisert EE, Lu L, Freeman-Anderson NE, et al. Gene expression in the mouse eye: an online resource for genetics using 103 strains of mice. Mol Vis 2009;15:1730–63.

15. King R, Struebing FL, Li Y, et al. Genomic locus modulating corneal thickness in the mouse identifies POU6F2 as a potential risk of developing glaucoma. PLoS Genet 2018;14(1):e1007145.

16. King R, Li Y, Wang J, et al. Genomic Locus Modulating IOP in the BXD RI Mouse Strains. G3 (Bethesda) 2018.

17. Struebing FL, King R, Li Y, et al. Genomic loci modulating retinal ganglion cell death following elevated IOP in the mouse. Exp Eye Res 2018;169:61–7.

18. Wang J, Geisert EE, Struebing FL. RNA sequencing profiling of the retina in C57BL/6J and DBA/2J mice: Enhancing the retinal microarray data sets from GeneNetwork. Mol Vis 2019;25:345–58.

19. Wang J, Li Y, King R, et al. Optic nerve regeneration in the mouse is a complex trait modulated by genetic background. Mol Vis 2018;24:174–86.

20. Bailey JN, Loomis SJ, Kang JH, et al. Genome-wide association analysis identifies TXNRD2, ATXN2 and FOXC1 as susceptibility loci for primary open-angle glaucoma. Nat Genet 2016;48(2):189–94.

21. Wiggs JL, Hauser MA, Abdrabou W, et al. The NEIGHBOR consortium primary open-angle glaucoma genome-wide association study: rationale, study design, and clinical variables. J Glaucoma 2013;22(7):517–25.

22. Springelkamp H, Mishra A, Hysi PG, et al. Meta-analysis of Genome-Wide Association Studies Identifies Novel Loci Associated With Optic Disc Morphology. Genet Epidemiol 2015;39(3):207–16.

23. Craig JE, Han X, Qassim A, et al. Multitrait analysis of glaucoma identifies new risk loci and enables polygenic prediction of disease susceptibility and progression. Nat Genet 2020.

24. Zhou H, Yoshioka T, Nathans J. Retina-derived POU-domain factor-1: a complex POU-domain gene implicated in the development of retinal ganglion and amacrine cells. J Neurosci 1996;16(7):2261–74.

25. Struebing FL, Lee RK, Williams RW, Geisert EE. Genetic Networks in Mouse Retinal Ganglion Cells. Front Genet 2016;7:169.

26. Groman-Lupa S, Adewumi J, Park KU, Brzezinski Iv JA. The Transcription Factor Prdm16 Marks a Single Retinal Ganglion Cell Subtype in the Mouse Retina. Invest Ophthalmol Vis Sci 2017;58(12):5421–33.

27. Martersteck EM, Hirokawa KE, Evarts M, et al. Diverse Central Projection Patterns of Retinal Ganglion Cells. Cell Rep 2017;18(8):2058–72.

28. Samuel MA, Zhang Y, Meister M, Sanes JR. Age-related alterations in neurons of the mouse retina. J Neurosci 2011;31(44):16033–44.

29. Voigt T. Cholinergic amacrine cells in the rat retina. J Comp Neurol 1986;248(1):19–35.

30. Carpenter AE, Jones TR, Lamprecht MR, et al. CellProfiler: image analysis software for identifying and quantifying cell phenotypes. Genome Biol 2006;7(10):R100.

31. Lamprecht MR, Sabatini DM, Carpenter AE. CellProfiler: free, versatile software for automated biological image analysis. Biotechniques 2007;42(1):71–5.

32. Dordea AC, Bray MA, Allen K, et al. An open-source computational tool to automatically quantify immunolabeled retinal ganglion cells. Exp Eye Res 2016;147:50–6.

33. Kay JN, De la Huerta I, Kim IJ, et al. Retinal ganglion cells with distinct directional preferences differ in molecular identity, structure, and central projections. J Neurosci 2011;31(21):7753–62.

34. Ivanova E, Lee P, Pan ZH. Characterization of multiple bistratified retinal ganglion cells in a purkinje cell protein 2-Cre transgenic mouse line. J Comp Neurol 2013;521(9):2165–80.

35. Sanes JR, Masland RH. The types of retinal ganglion cells: current status and implications for neuronal classification. Annu Rev Neurosci 2015;38:221–46.

36. De la Huerta I, Kim IJ, Voinescu PE, Sanes JR. Direction-selective retinal ganglion cells arise from molecularly specified multipotential progenitors. Proc Natl Acad Sci U S A 2012;109(43):17663–8.

37. Sweeney NT, James KN, Nistorica A, et al. Expression of transcription factors divides retinal ganglion cells into distinct classes. J Comp Neurol 2019;527(1):225–35.

38. Dhande OS, Stafford BK, Franke K, et al. Molecular Fingerprinting of On-Off Direction-Selective Retinal Ganglion Cells Across Species and Relevance to Primate Visual Circuits. J Neurosci 2019;39(1):78–95.

39. Macosko EZ, Basu A, Satija R, et al. Highly Parallel Genome-wide Expression Profiling of Individual Cells Using Nanoliter Droplets. Cell 2015;161(5):1202–14.

40. Clark BS, Stein-O’Brien GL, Shiau F, et al. Single-Cell RNA-Seq Analysis of Retinal Development Identifies NFI Factors as Regulating Mitotic Exit and Late-Born Cell Specification. Neuron 2019;102(6):1111–26 e5.

41. Rheaume BA, Jereen A, Bolisetty M, et al. Single cell transcriptome profiling of retinal ganglion cells identifies cellular subtypes. Nat Commun 2018;9(1):2759.

42. Tran NM, Shekhar K, Whitney IE, et al. Single-Cell Profiles of Retinal Ganglion Cells Differing in Resilience to Injury Reveal Neuroprotective Genes. Neuron 2019;104(6):1039–55 e12.

43. Bray ER, Noga M, Thakor K, et al. 3D Visualization of Individual Regenerating Retinal Ganglion Cell Axons Reveals Surprisingly Complex Growth Paths. eNeuro 2017;4(4).

44. Duan X, Qiao M, Bei F, et al. Subtype-specific regeneration of retinal ganglion cells following axotomy: effects of osteopontin and mTOR signaling. Neuron 2015;85(6):1244–56.

45. Daniel S, Clark AF, McDowell CM. Subtype-specific response of retinal ganglion cells to optic nerve crush. Cell Death Discov 2018;4:7.

